# Adenovirus type 34 and HVR1-deleted Adenovirus type 5 do not bind to PF4: clearing the path towards vectors without thrombosis risk

**DOI:** 10.1101/2022.11.07.515483

**Authors:** Erwan Sallard, Daniel Pembaur, Katrin Schröer, Sebastian Schellhorn, Georgia Koukou, Natascha Schmidt, Wenli Zhang, Florian Kreppel, Anja Ehrhardt

## Abstract

The adenoviral vector based AstraZeneca and Janssen COVID vaccines have been associated with rare cases of thrombosis, a condition which depends on adenovirus binding to the blood protein Platelet Factor 4 (PF4).

In order to identify adenoviruses with low or absent affinity for PF4, we screened dozens of types from various adenovirus species, and Adenovirus type 5 (Ad5) derived vectors carrying genetic or chemical modifications of different hexon hyper-variable regions (HVR). For this purpose, we established an armamentarium of techniques including ELISA-qPCR and Aggregate Pull-Down (APD), which enabled fast and sensitive assessments of virus-protein interactions.

Unlike most tested serotypes, Ad34 did not bind to PF4. Likewise, the deletion or shielding of the HVR1 loop of Ad5 seemingly ablated its PF4 binding. Therefore, we showed that PF4 binds to adenovirus hexon through interactions dependent on HVR1, and identified vectors that may avoid or decrease the risk of thrombosis and represent safer candidates for vaccine or gene therapy vector development.

## Introduction

Adenoviruses (Ads) are non-enveloped viruses with a linear double-stranded DNA genome comprising between 26 and 45 kb (*1*). There are currently 113 known Ad types infecting humans (*2*), in addition to an even larger diversity of non-human Ads infecting other species including primates. Thanks to their high manufacturability, gene delivery efficiency, genetic stability and ability to package large transgenes, Ad-derived vectors are the most prominent type of vectors used in gene therapy or vaccine development (*3, 4*). Notably, the AstraZeneca (ChAdOx1-nCoV19, derived from chimpanzee Ad type Y25, hereby termed ChAdY25) and Janssen (derived from human Ad26, hereby designated after its capsid as Ad26) COVID vaccines have already been administered well over 2 billion times (*5*), and established Ad vaccines as one of the most powerful tools against pandemics.

However, clinical applications of Ad vectors still face several obstacles, among which the ability of certain Ad types to interact with blood proteins after systemic administration or local injection, including with prothrombin, the most abundant coagulation factor (*6*). Moreover, Ad type 5 (Ad5) displays a strong and potentially pathological liver tropism due to its binding to the coagulation factor X (fX) (*7*) on the fifth and seventh hypervariable regions (HVR) of its hexon protein (*8*), the main Ad capsid protein. Likewise, the ChAdY25 and Ad26 vaccines have been associated with very rare cases of vaccine-induced immune thrombotic thrombocytopenia (VITT), with an incidence in the order of magnitude of 1 case per 100,000 vaccinated persons (*9*). This disorder usually occurs within 5 to 20 days after the first vaccine injection, involves thrombosis in the cerebral venous sinus, the splanchnic vein or other unusual thromboembolic events (*10*), and is lethal in 23-40% of cases (*11*, *12*). It was recently discovered that VITT is caused by the binding of the vectors to platelet factor 4 (*10*, *13*) (PF4, also known as CXCL4), which can activate a cascade of immune reactions, notably the production of auto-antibodies, leading to severe adverse effects in a small subset of patients. PF4 is a 7.8 kDa cationic protein secreted by activated thrombocytes whose physiological function is thought to be the recruitment of thrombocytes on glycosaminoglycans exposed in vascular injuries and / or the opsonization of the negatively charged surfaces of pathogens (*14*). PF4 blood concentrations lies usually around 10 ng/mL, but can reach 3-15 μg/mL in case of platelet activation (*15*, *16*).

The development of safer vaccine and gene therapy vector platforms may protect patients from rare but fatal side effects and improve public trust in medical treatments and prophylaxes. Therefore, we aimed to identify Ad types with lower PF4 binding. We screened a collection of 23 reporter gene carrying Ad vectors from different types, representative of the natural diversity of human-infecting Ads (*17*). To this purpose, we established a variety of on-target techniques for fast, robust and sensitive assessment of protein-virus interactions. Our techniques revealed that Ad34 lacked detectable PF4 binding. Furthermore, using a series of Ad5 variants carrying genetic or chemical hexon modifications, we found that HVR1 deletion or shielding also ablated PF4 binding. This confirmed the hypothesis that PF4 binds adenoviruses on the hexon protein, and identifies the HVR1 hexon loop as one critical interaction site.

## Results

### Screening of an Ad collection indicated that Ad34 lacked PF4 binding

In order to rapidly screen a large Ad collection, we established the ELISA-qPCR technique (figure 1a, supplementary figure 1) with which we could confirm preexisting data (*13*) by detecting PF4 binding of Ad5, ChAdY25 and Ad26 (figure 1b).

**Figure 1:**
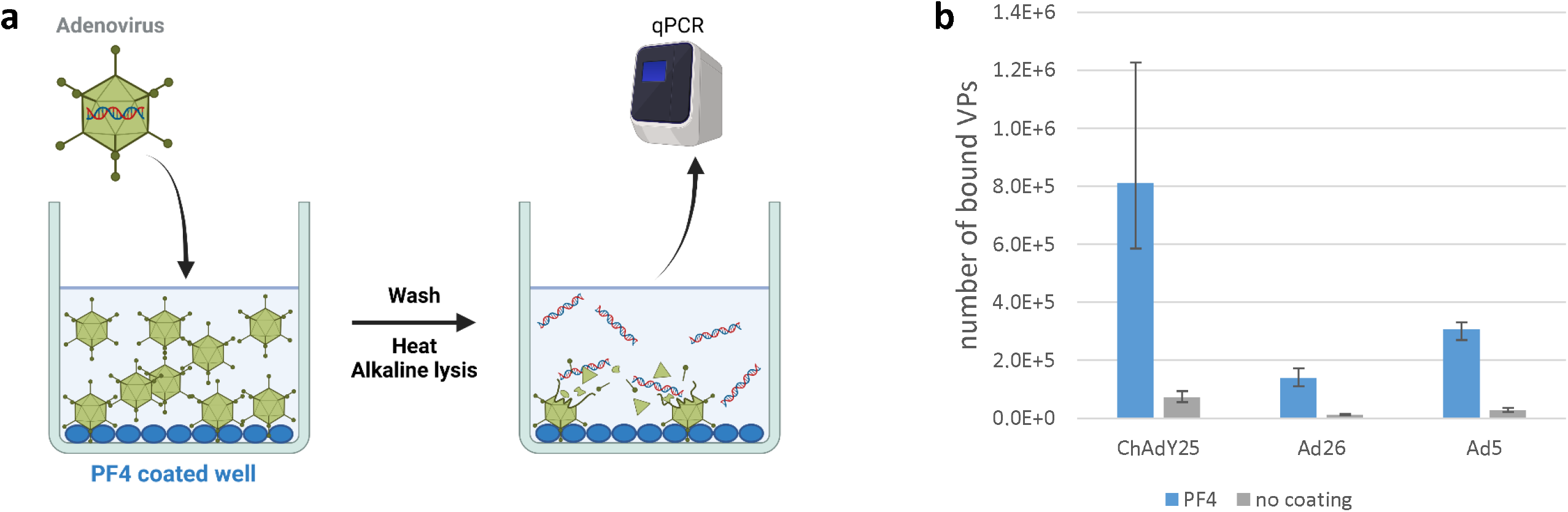
The novel ELISA-qPCR technique facilitates the detection of PF4-Adenovirus interactions. **(a)** Principle of the ELISA-qPCR technique. Adenovirus (Ad) VPs are allowed to interact with proteins, e.g. PF4, coated on an ELISA plate. After washes, the genomes of VPs which specifically interacted with the proteins are released by heating and alkaline treatment and quantified by qPCR. Figure created with BioRender. (**b)** PF4 binding of vaccine-equivalent vectors. Ad5 was obtained from the Ad-GLN library. N=4, two independent repeats.

Using ELISA-qPCR, we screened 23 different human Ad types for PF4 binding. Ad34 and Ad50 were the only types for which PF4 binding could not be detected in any of the experimental repeats (figure 2a, supplementary figure 2).

**Figure 2:**
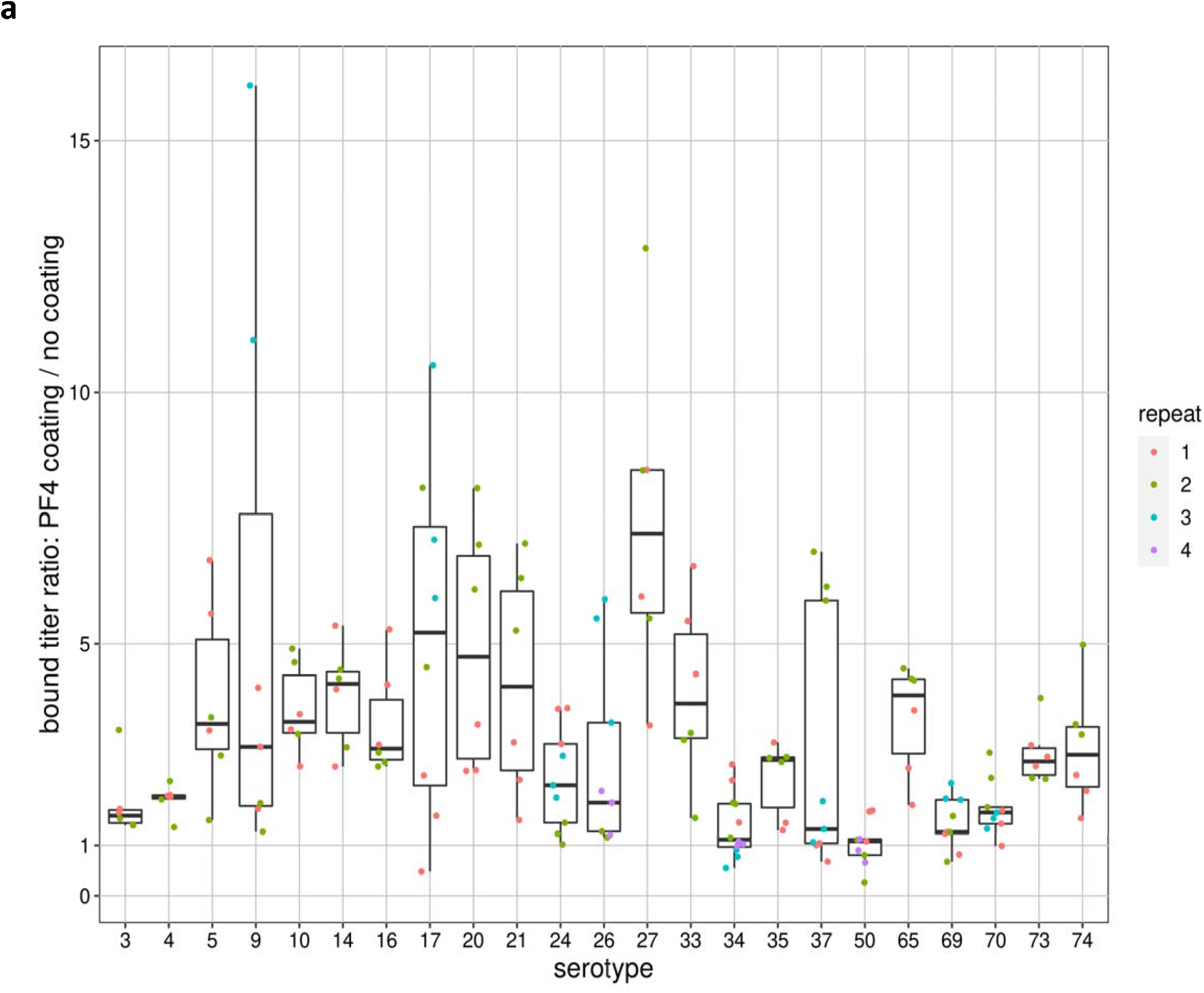
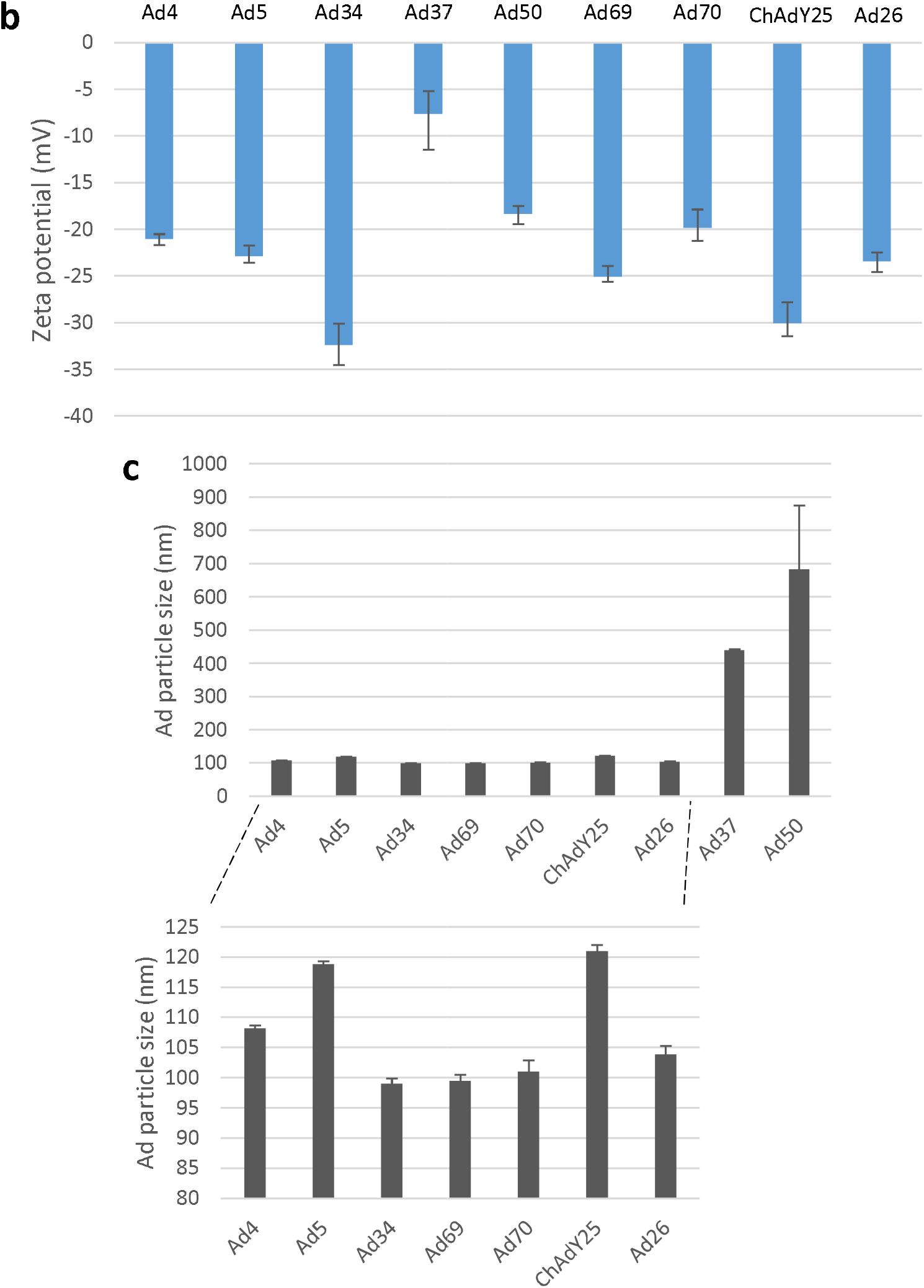
Ad34 does not bind to PF4. **(a)** Screening of the Ad-GLN library for PF4 binding by ELISA-qPCR. Each point represents the number of bound VPs in one PF4 coated well normalized on the average number from the three non-coated wells of the same experiment repeat. N≥6, two to four independent repeats. **b, c:** Surface potential **(b)** and hydrodynamic size **(c)** of Ad particles measured by dynamic light scattering (DLS). Ad types with high PF4 binding (Ad5, ChAdY25, Ad26), medium to low binding (Ad4, Ad69, Ad70, Ad37) as well as no binding (Ad34, Ad50) according to ELISA-qPCR assays were included. Sizes larger than the usual 80-130nm of Ad particles indicate agregate formation. ChAdY25, Ad26: vectors with capsids equivalent to AstraZeneca and Janssen vaccines respectively. The other vectors belong to the Ad-GLN library. N=3.

Since PF4 binding on Ads may be driven by electrostatic interactions between the negative hexon protein and the positive PF4 (*13*), we studied by electrophoretic light scattering (ELS) whether Ad binding strength to PF4 was correlated with a net negative surface charge measured as zeta potential. However, the more negative potential of Ad34 compared with other Ads showed that a negatively charged surface was not sufficient for PF4 binding (figure 2b). These measurements nevertheless indicated that our Ad50 suspension contained viral particle (VP) aggregates and its apparent lack of PF4 binding may be an artifact (figure 2c).

### PF4 increased aggregate formation, cell binding and infection of permissive cells by Ad5 and Ad69 but not Ad34

We sought to confirm the results obtained so far by screening with independent techniques a subset of vectors which had shown varying PF4 binding strength, namely Ad5, ChAdY25 (strong binding), Ad69 (weak binding), and Ad34 (no detectable binding). First, we used the novel Aggregate Pull-Down (APD) technique to quantify Ad VP aggregates that may form upon interaction with PF4 (figure 3a). As expected, we observed PF4-induced aggregation of Ad5, ChAdY25 and Ad69, but not Ad34, confirming the lack of PF4 binding of this Ad type (figure 3b).

**Figure 3:**
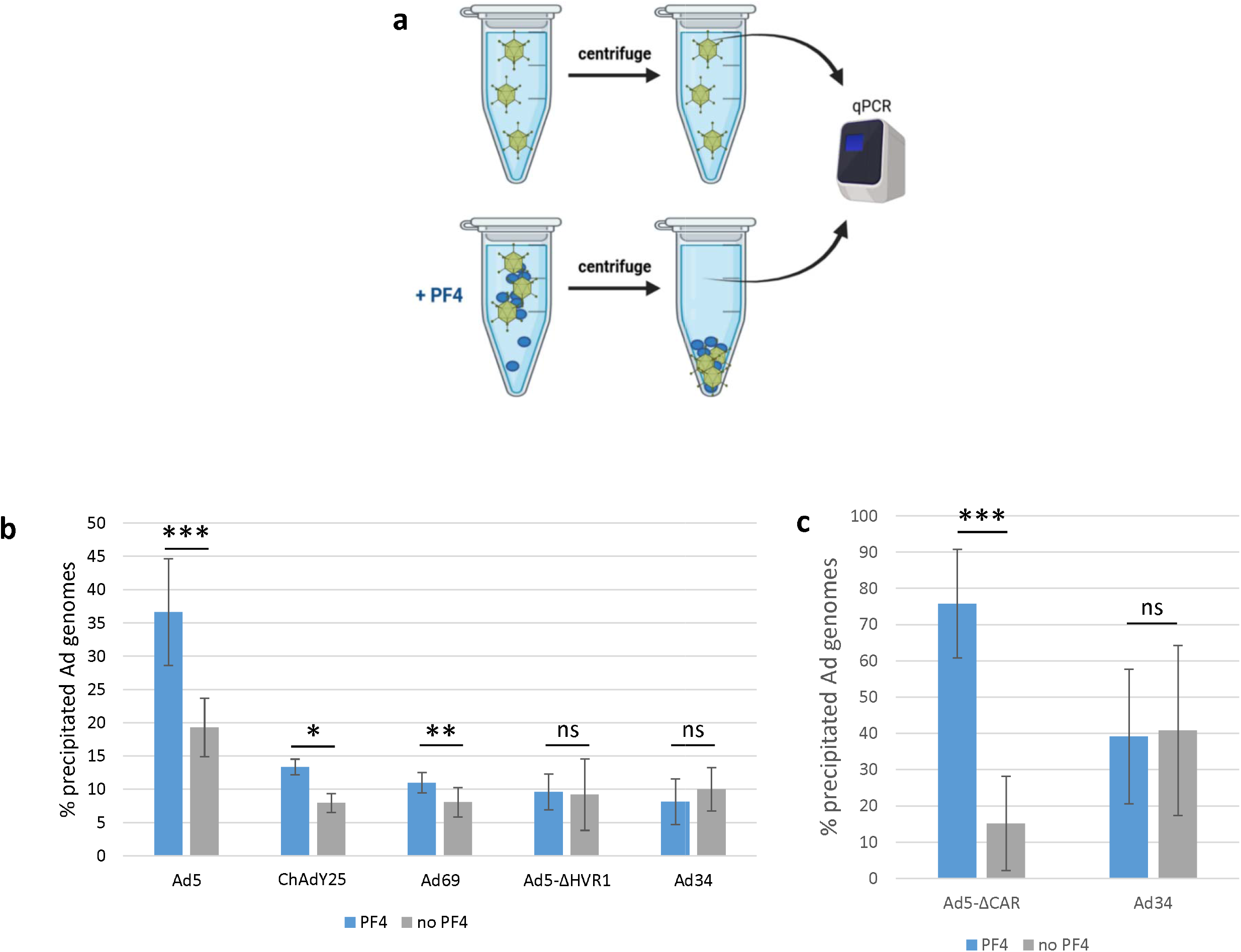
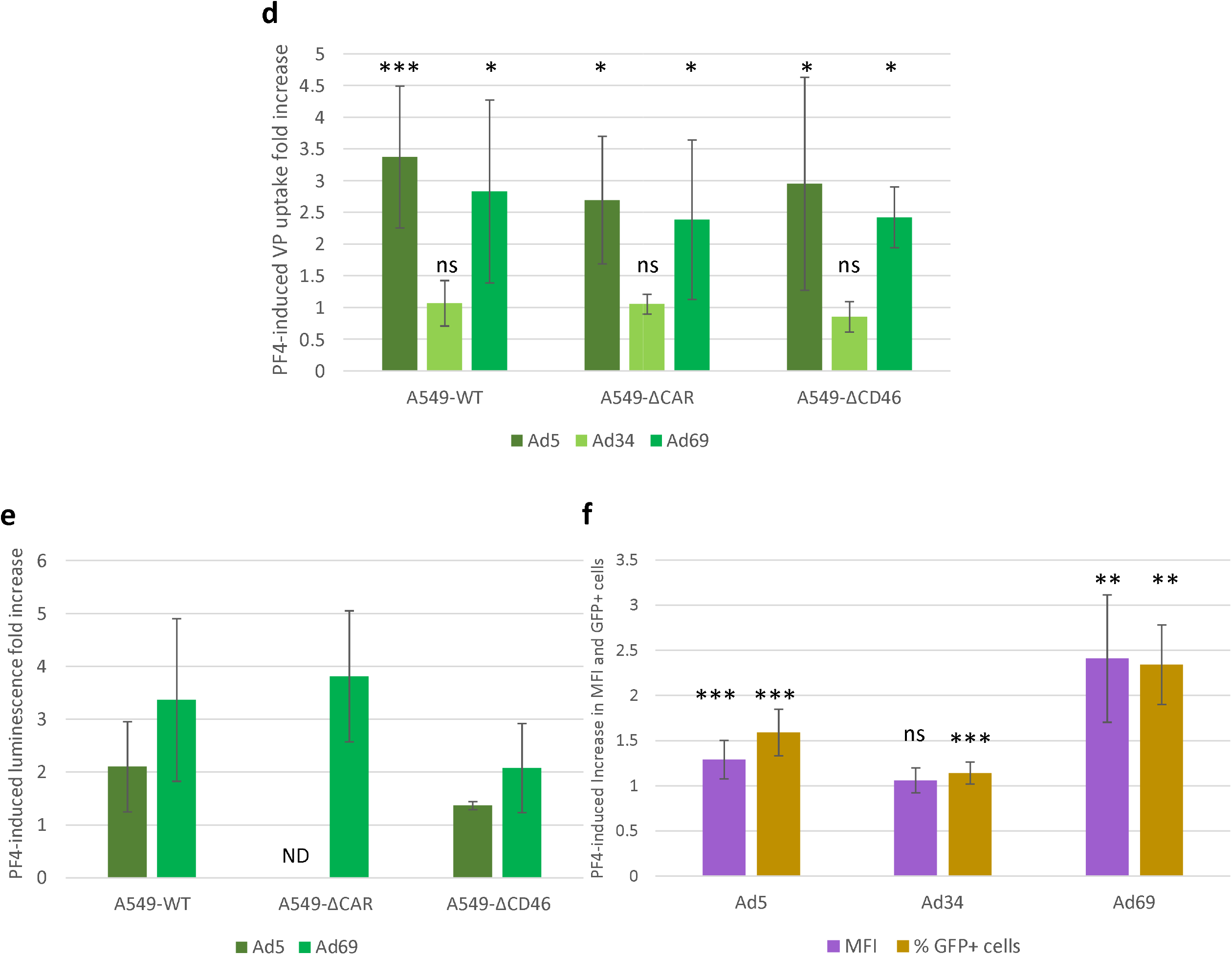
PF4 increases aggregation, cell binding and infectivity of Ad5 and Ad69 but not Ad34. **(a)** Principle of the Aggregate Pull-Down technique. Aggregates forming upon interaction with PF4 are separated from free VPs by low speed centrifugation and titrated by qPCR. Figure created with BioRender. **(b)** Aggregate pull-down of selected Ads in absence or presence of PF4. Ad5, Ad69 and Ad34 were obtained from the Ad-GLN library. N=8, two independent repeats. Pairwise comparisons were conducted for each Ad with the Mann-Whitney U test. **(c)** Erythrocyte pull-down of selected Ads in absence or presence of PF4. We used a fiber-modified Ad5 with ablated CAR tropism (Ad5-ΔCAR) and Ad34 from the Ad-GLN library. N≥10, three independent repeats. Pairwise comparisons were conducted for each Ad with the Mann-Whitney U test. **(d)** Fold change in Ad genome internalization in A549-derived cells following Ad incubation with PF4. VPs were incubated 10 min. at 37°C in optiMEM with or without 10μg/mL of PF4, before being allowed to infect cells at 20 VP/cell (vpc). 3 hours post infection (hpi), internalized Ad genomes were titrated by qPCR. A549-ΔCAR cells lack the primary receptor of Ad5 and Ad69, while A549-ΔCD46 lack the primary receptor of Ad34. All Ads belong to the GLN library. N≥6, three to five independent repeats. Wilcoxon signed ranks tests were used to assess whether PF4-associated fold increases are significantly different from 1. Welch’s ANOVA did not detect significant effect of the cell line on PF4-associated fold increase (p = 0.47, 0.32 and 0.81 for Ad5, Ad34 and Ad69 respectively), **(e)** Fold change in luciferase luminescence in A549-derived cells following Ad incubation with PF4. VPs were incubated 10 min. at 37°C in optiMEM with or without 10μg/mL of PF4, before being allowed to infect cells at 20 vpc. 24 hpi, the ratio of luciferase luminescence (arbitrary units) between samples with PF4 co-incubation and those without was measured. All Ads belong to the GLN library. N≥4, two to three independent repeats. **(f)** Infectivity assay with fluorescence readout. VPs were incubated 10 min. at 37°C in optiMEM with or without 10μg/r⋂L of PF4, before being allowed to infect cells at 20 vpc. 24 hpi, Ad-expressed GFP mean fluorescence intensity (MFI) and proportion of GFP-positive cells (% GFP+ cells) were measured by flow cytometry. Ad34 and Ad69 were obtained from the Ad-GLN library, while an El-deleted, GFP-expressing Ad5 vector was used. N≥9, three to four independent repeats. Wilcoxon signed ranks tests were used to assess whether PF4-associated fold increases are significantly different from 1.

Ads are able to bind to erythrocytes (*18*), leading to substantial vector sequestration and retargeting following systemic administrations. Therefore, studying the influence of Ad-PF4 interactions on erythrocyte binding may yield interesting first hints into their *in vivo* consequences. We observed that PF4 had no influence on Ad34 but substantially increased erythrocyte docking of a fiber-modified Ad5 with ablated CAR tropism (Ad5-ΔCAR, figure 3c).

We finally tested whether PF4 binding impacted Ad infectivity. Incubation of Ad5 or Ad69 particles with PF4 increased VP uptake in A549 cells by around 2 to 3 fold in average, while PF4 did not influence Ad34 internalization (figure 3d). PF4 effect on infectivity was not significantly altered by deletion of CAR (primary receptor of Ad5 and Ad69 (*19*)) or CD46 (primary receptor of Ad34 and Ad69 (*19*)) in target cells (figure 3d). In order to test whether PF4 enhanced productive infections by Ad5 and Ad69 or only abortive internalizations, we measured Ad-driven luciferase expression in infected cells and detected an increase for both Ads and in all tested cell lines (figure 3e). Likewise, PF4 increased Ad-driven GFP expression in cells infected by Ad5 and Ad69 (figure 3f). For Ad34, PF4 did not modify the mean GFP fluorescence intensity but was associated with a 14% increase in the proportion of GFP-positive cells (figure 3f).

### The hexon HVR1 loop is necessary for Ad5 binding of PF4

Baker *et al*. predicted by brownian dynamics modeling that PF4 binds to hexon HVRs (*13*), with HVR1 being the most likely candidate for the ChAdY25 vaccine. In order to gather further information on the location of PF4 binding site(s), we compared by ELISA-qPCR the PF4 binding of Ad5 variants with chemically or genetically modified hexons (figure 4a). Point mutations in HVR1, HVR5 and HVR7, the latter being known to ablate fX binding (*20*), did not prevent PF4 binding, contrary to deletion of the full HVR1 loop. PEGylation of HVR1 and HVR5, i.e. covalent linking of a large inert polymer which sterically prevents interactors to bind near its linkage site, also inhibited PF4 binding (figure 4b). The lack of PF4 binding by the HVR1-deleted Ad5 was confirmed by APD (figure 3b).

**Figure 4:**
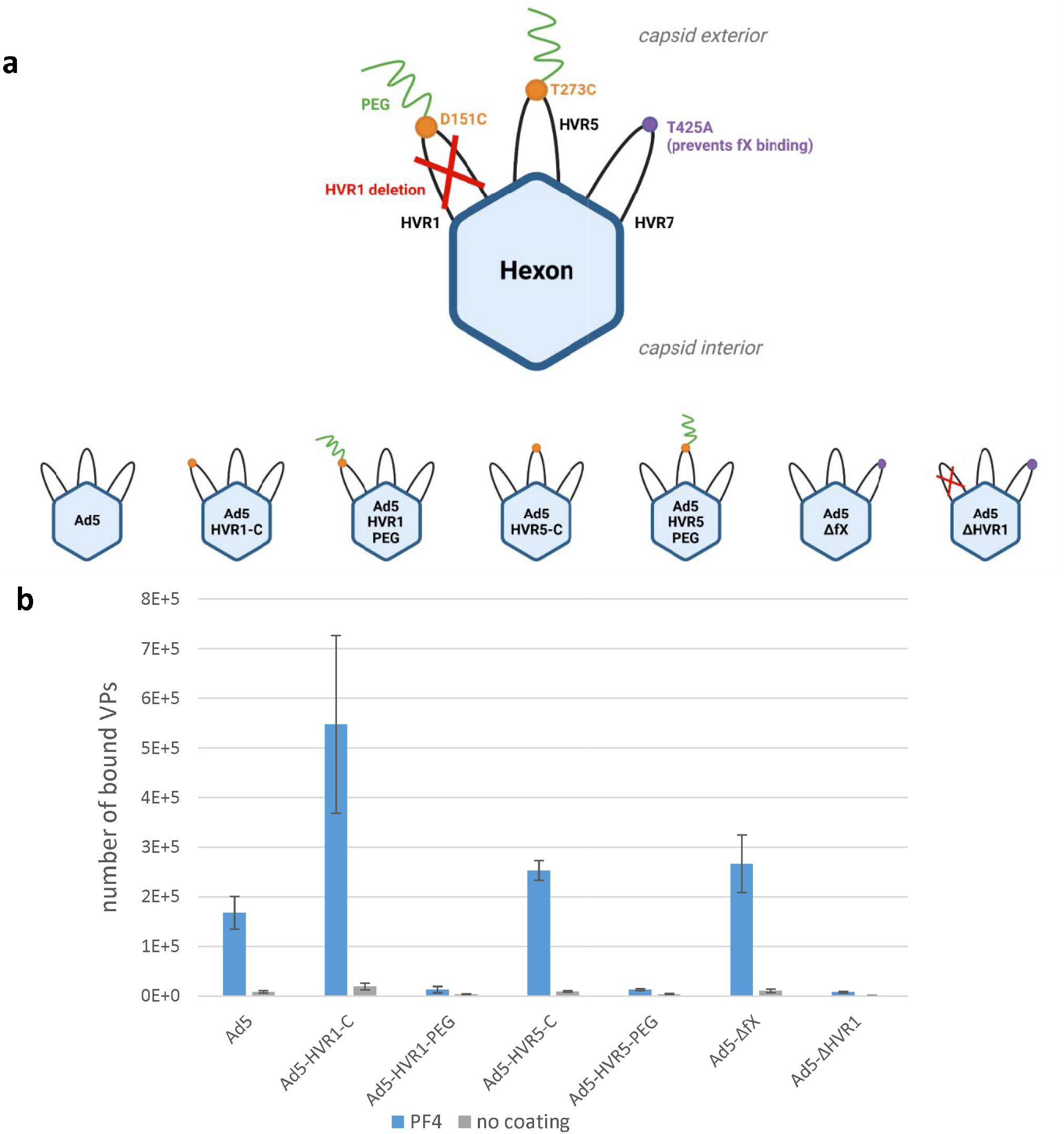
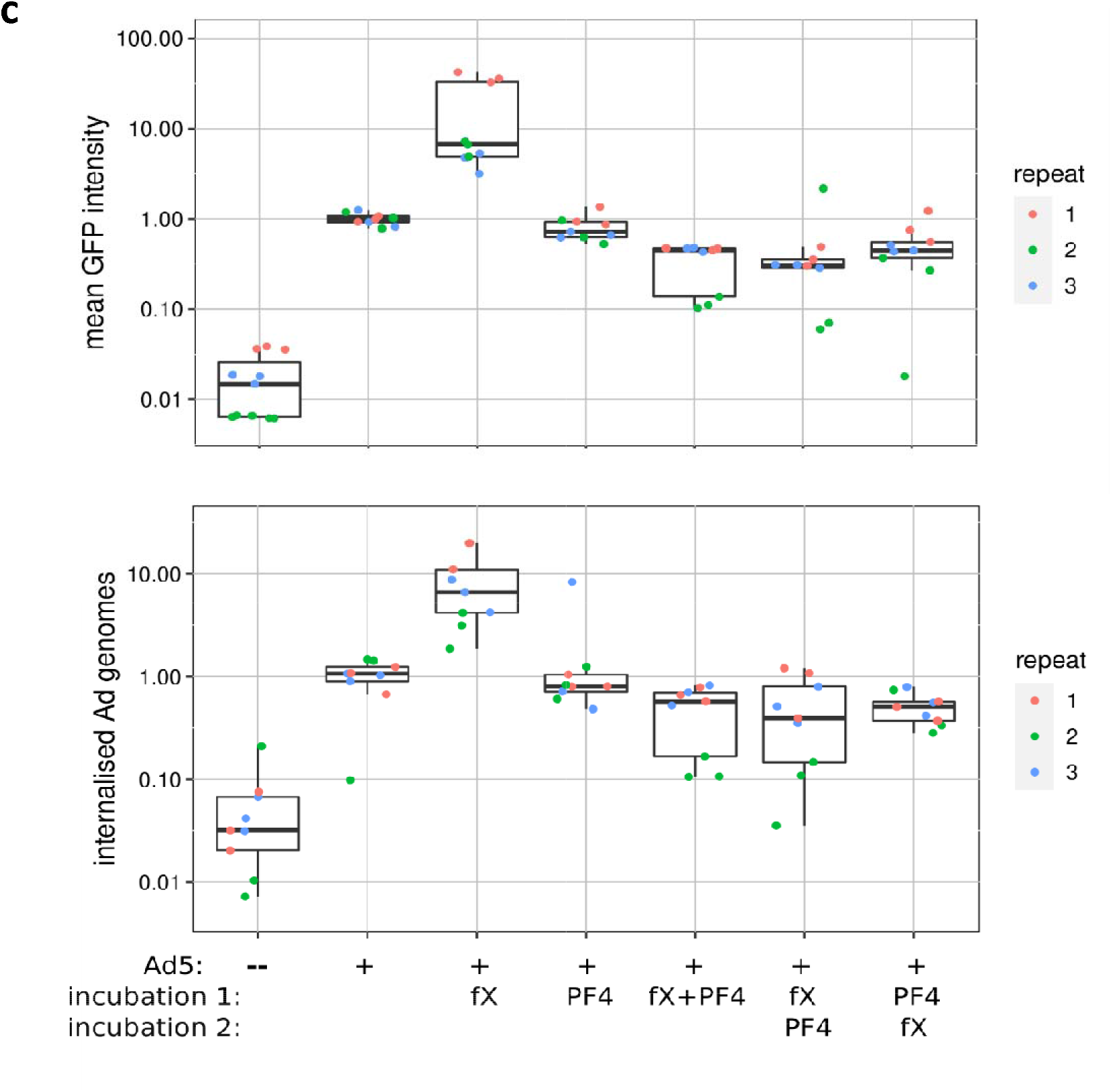
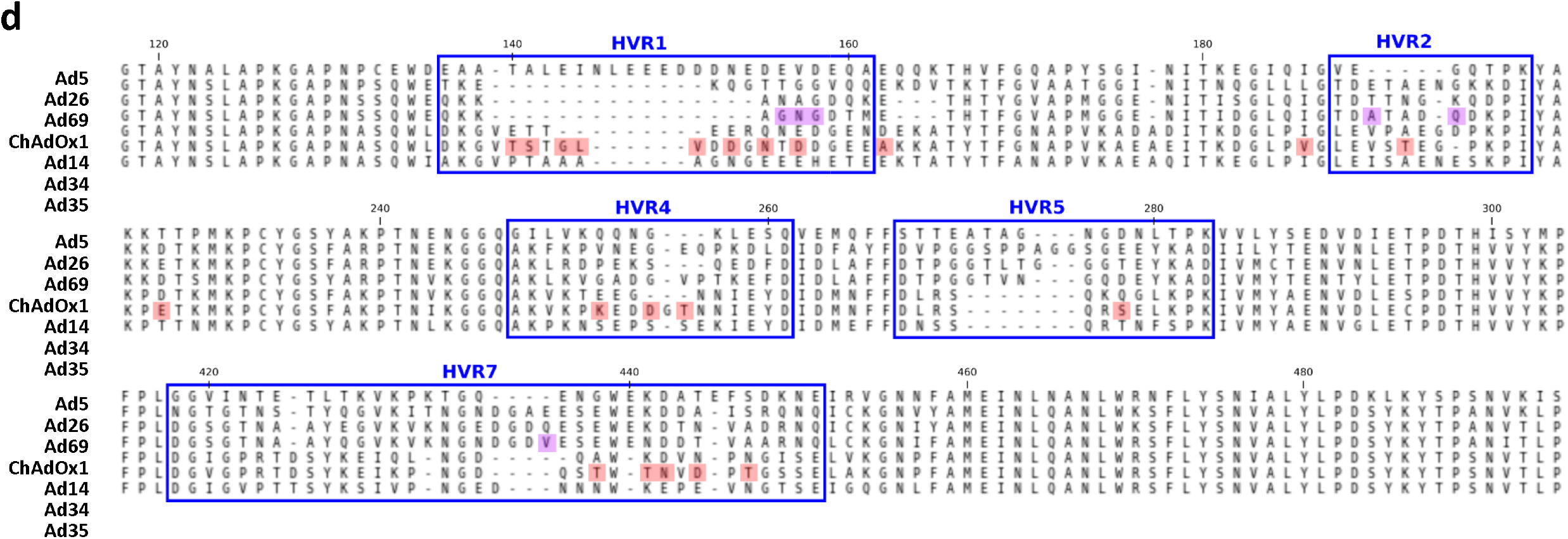
PF4 do not compete with factor X for binding on Ad5 hexon. **(a)** Schematic representation of the Ad5 hexon genetic and chemical variants studied. These variants include: D151C and T273C point mutations; covalently linking a cysteine residue with a 5kDa polyethylene-glycol (PEG) polymer, which prevents binding on part of the hexon surface by steric competition; deletion of the HVR1 loop; and T425A substitution, which ablates the binding of fX. The El-deleted, GFP-expressing Ad5 vector was used as control (Ad5). HVR: hyper-variable region. PEG: poly-ethylene glycol. Figure created with BioRender, **(b)** ELISA-qPCR of the Ad5 hexon variants for PF4 binding. N=6, two independent repeats, **(c)** PF4 inhibits fX-dependent Ad5 infectivity of SKOV-3 cells. The El-deleted, GFP-expressing Ad5 vector was incubated (from left to right) in optiMEM, with fX, PF4, or both proteins simultaneously or fX first or PF4 first. VPs were then allowed to infect SKOV-3 cells. Infectivity was measured either by quantification of GFP fluorescence intensity by flow cytometry 3 days post infection (top) or qPCR titration of internalized genomes 3 hours post infection (bottom). qPCR genome copy numbers and fluorescence intensity were normalized on the average of the, Ad5 alone” sample. N=9, three independent repeats. For both measurement types, pairwise comparisons were conducted by Mann-Whitney U test between the „Ad5 + fX” sample and each of the other samples, always indicating significant difference with p<0.001. **(d)** Hexon residues specific of Ad34 cluster in HVR loops. The hexon sequences of serotypes with strong (Ad5, Ad14, Ad26, ChAdOx), weak (Ad35, Ad69) or no (Ad34) PF4 binding were aligned with the MAFFT online tool. HVRs are indicated by blue rectangles, residues specific to Ad34 among the chosen serotypes in red, and ChAdOx1 residues with high or medium probability of interaction with PF4 according to Baker *et al*. modelisation in purple. Numbers above the sequences indicate the amino acid position in Ad5 hexon sequence.

SKOV-3 cells are permissive to Ad5-fX complexes, but largely refractory to free Ad5 VPs (*21*). We observed that Ad5 incubation with PF4 prior, simultaneously or following incubation with fX inhibited fX-driven SKOV-3 cell infection (figure 4c). Since this phenomenon occurred irrespective of the incubation order, this may suggest that there is no competition between PF4 and fX for binding on Ads.

We compared the hexon structure and sequence of Ad34 with other Ad types, including the closely related (*22*) but PF4-binding Ad35 and Ad14, in order to identify structural elements unique to Ad34 which may explain its apparent lack of PF4 binding. A large proportion of residues unique to Ad34 are clustered in HVR1 (figure 4d), corresponding to the most likely binding site candidate identified by Baker *et al*. HVR1 is predicted to be the only hexon site with substantially different conformation in Ad34 compared with Ad35 and Ad14 (supplementary video 1).

## Discussion

Here we establish an armamentarium composed of user-friendly, scalable and affordable techniques for the study of virus–protein interactions including ELISA-qPCR, APD and infectivity assays. We confirmed the specificity of ELISA-qPCR in the case of a few known interactors of Ad5 (supplementary figure 1a). This method also displayed a relatively high sensitivity, as shown by the significant detection of PF4-Ad5 binding even at low VP concentration and despite the relatively low affinity of this interaction (K_D_ = 789 nM (*13*)). Our assays almost systematically yielded concordant results and constitute a panoply of independent techniques facilitating the screen of large Ad collections for PF4 binding with results robust across experimental conditions. However, our assays remain so far only qualitative and appeared sensitive to the quality of virus preparation (figure 2c, supplementary figure 2). Further experiments using varying concentrations of VPs and proteins are warranted to elucidate their sensitivity cut-offs. Finally, additional tests should assess the range of applications of our new techniques, which in theory extend to all virus-protein interactions in the case of ELISA-qPCR.

We started to investigate the consequences of PF4 binding on Ad tropism and potential *in vivo* interactions. PF4 induced a strong increase in erythrocyte binding of a fiber-modified Ad5 vector (figure 3c). Likewise, PF4 induced a strong increase in Ad5 and Ad69 infectivity of permissive cells (figure 3d,e,f). Measurements of Ad5 luciferase expression (at 24 hours post infection) indicated a lower PF4-induced infectivity increase than VP uptake measurements (at 3 hours post infection), and could not be conducted in A549-ΔCAR cells, because of Ad5 toxicity (figure 3e). We therefore used a replication-deficient Ad5 vector with a WT capsid for our GFP fluorescence measurements (figures 3f, 4c). For both Ad5 and Ad69, a similar increase in infectivity was observed in cells lacking the virus primary receptors, suggesting that the infectivity increase does not result from specific interactions between PF4 and a cellular receptor. We hypothesize that it instead stems from a partial neutralization of Ad particle surface, which decreases electrostatic repulsion with the extracellular matrix, similar to cationic polymers used to improve the delivery of non-viral particles (*23–25*). In order to test if the PF4-mediated infectivity increase is explained only by the formation of VP aggregates (as observed by APD) or if PF4 also increased the infectivity of free VPs, we also quantified Ad-driven GFP fluorescence in infected A549 cells (figure 3f). Since not only the mean fluorescence intensity (MFI) but also the proportion of GFP-positive cells was increased by preincubation of Ad5 and Ad69 (but once again not Ad34) with PF4, we favour the second hypothesis.

In the same experiment, PF4 surprisingly appeared to induce a small but significant increase in the percentage of GFP-positive cells after Ad34 infection. This contradicted all other results regarding Ad34, which until then had never been affected by the presence of PF4. This may stem from an influence of PF4 on cell viability or metabolism affecting GFP expression, decrease in virus samples quality over time, or indicate a very weak PF4 binding by Ad34 that remained under the detection threshold of the other methods used. In our opinion, this does not exclude Ad34 as a promising candidate for vectors with reduced PF4 binding, but warrants testing its PF4 affinity with more accurate methods such as SPR.

The inhibition of PF4 binding by HVR1 deletion (figure 4b) and PEGylation of HVR1 and HVR5 (which is spatially very close to HVR1) proved that PF4 binds to the hexon of Ad5. Moreover, these results and the comparison of hexon structures and sequences of PF4 binder versus non-binder Ad types (figure 4d) point to HVR1 as the most likely binding site. Furthermore, the inhibition of fX-dependent SKOV-3 infection by Ad5 after coincubation with PF4 and fX (figure 4c) suggested that PF4 inhibits an infection step downstream of fX. Indeed, the affinity of Ad5 for PF4 is substantially lower than its affinity for fX (K_D_ = 789 nM (*13*) versus 1.83 nM (*7*) or 2.7 nM (*26*)), thus fX would probably prevail over PF4 if a competition occurred, even though other factors such as PF4-fX interactions may be at play. Combined with the observation that mutations that ablate fX binding had no impact on PF4 binding (figure 4b), this suggested that the PF4 binding site is separated from that of fX. However, the study of additional hexon variants would be required to fully elucidate the binding site of PF4 and whether it interferes with fX for hexon docking. Furthermore, given the flexibility of hexon HVRs, it can be envisioned that PF4 does not have a unique binding site.

Like other *in vitro* studies, our investigation of PF4 binding to Ads either in optiMEM, PBS or immobilized on an ELISA plate may not fully recapitulate the interactions occurring in the blood or in tissues, which are more complex environments in which other interaction partners can be involved. In particular, understanding the immunological consequences of the modified Ad cell binding and infection patterns observed here will require dedicated investigation. Still, we can reasonably expect that the Ad types for which we detected no PF4 binding would at least display lower binding *in vivo* than other Ads.

Adenoviral vaccines, gene therapy vectors and oncolytic vectors without VITT risk may prevent a number of patients deaths and even more non-fatal complications, as well as improve the public trust in medical treatments and adherence epidemics containment measures. Since we have not observed conclusive PF4 binding to Ad34, using several techniques and three different virus preparations of confirmed quality, this type should be considered as a promising vector candidate. Adding to its potential high safety profile, Ad34 has a very low seroprevalence (Wang *et al*., Journal of Virology, in press). Alternatively, the safety of existing vaccine platforms may be directly improved by HVR1 deletion or pseudotyping with Ad34 HVR1. Moreover, Ad types not included in this study, notably simian Ads, may be screened for PF4 binding using the panoply of techniques established in this article (ELISA-qPCR, infectivity assays and Ad-PF4 aggregates or erythrocytes pull-down) in order to identify novel candidates and gather new insights about Ad-blood interactions and VITT.

Our data unequivocally demonstrated that polymer shielding is a means to control the unwanted interaction with PF4. Future developments on polymer modification of non-Ad5 types will be required to assess its full potential, notably in the context of repeated Ad delivery as often required by vaccination or therapy regimens.

Thrombotic disorders have been associated not only with adenoviral vaccines, but also adeno-associated gene therapy vectors (AAVs) (*27*). Even though the AAV-associated thrombotic microangiopathy differs from the Ad-related VITT in its clinical presentation, it is striking that the AAV9 type associated with all identified cases binds to PF4 (*28*). Thus, investigations on PF4 binding of non-Ad viral vectors and the potential pathologic consequences are warranted.

To conclude, we establish here a novel technique armamentarium facilitating fast, affordable, sensitive and specific assessment of virus-protein interactions. Furthermore, we identified several Ads, most notably Ad34 but also Ad5 hexon variants such as an HVR1-deleted mutant, which lack PF4 binding. These results may represent a milestone in the development of safer Ad vectors.

## Materials and Methods

### Vector acquisition

The Ad-GLN library, already described in (*17*), contains vectors from different types that express TurboGFP, NanoLuc luciferase and the selection marker kanamycin/neomycin under a synthetic CAG promoter in the deleted E3 region. The CAG promoter consists of the human cytomegalovirus early enhancer element, the chicken beta-actin promoter including parts of the first exon and intron, and parts of the second intron and third exon of the rabbit beta-globin gene.

The hexon-modified Ad5 vectors were produced similarly as described in (*29*). Briefly, genetic capsid modifications were introduced using pRed/ET homologous recombination (Gene Bridges, Heidelberg, Germany) in a bacmid carrying an Ad5 genome (AY339865) with the E1 locus (bp 441-3534) replaced by a CMV-promoter driven eGFP expression cassette. Vector PEGylation was conducted with 5 kDa mono-activated maleimide polyethylene glycol as described in (*29*).

The CAR-ablated Ad5 (described in (*30*)) was generated by introducing the point mutation Y477A (AAQ19310.1) in the fiber gene of the GFP-expressing, E1-deleted Ad5 backbone with the pRed/ET recombination kit.

The ChAdY25 and Ad26 vectors with capsids equivalent to the COVID vaccines were grown on HEK293 cells and purified by double CsCl banding and subsequent desalting with PD-10 columns (SE Healthcare).

### ELISA-qPCR

Proteins used in this study were human platelet factor 4 (PF4-h, Chromatec), which was stored at 4°C in PBS at a concentration of 200 μg/mL; human factor X (fX, Cellsystems #HCX-0050-MG); and S.typhimurium tRNA-specific adenosine deaminase (tadA, MyBioSource #MBS1445221). Proteins of interest were diluted in coating buffer (0.1 M NaHCO_3_, pH set between 9.2 and 9.6 using 1 M Na_2_CO_3_) to a concentration of 20 μg/mL and 75 μL were added per well of ELISA plate (Sarstedt #82.1581.200), which was sealed with a transparent film and incubated overnight at 4°C. Wells were washed twice with TBS-Tween (TBST), blocked with blocking buffer (TBST + 0.5% pork skin gelatin) 1 h at room temperature (RT), and washed twice with TBST. Virus particles were diluted in blocking buffer and incubated in the chosen coated wells for 2 h at 37°C. Except stated otherwise, 5E7 VP were used per well with 75 μL total volume. After virus incubation, wells were washed four times with TBST in order to eliminate VPs which did not interact specifically with the coated proteins. To quantify the remaining VPs, 50 μL of alkaline lysis buffer (25 mM NaOH + 2 mM EDTA) were added per well and the plate was carefully and tightly sealed and heated at 95°C for 10 min. to open capsids and release vector genomes. The plate was then immediately put on ice and 17 μL of cold neutralisation solution (80 mM Tris-Hcl + 0.1% Tween20, pH = 3.2) were added in each well. The virus genome solutions of each well were homogeneized by shaking and two 2 μL aliquots were taken for qPCR titration using the my-Budget 5x EvaGreen qPCR-Mix II (Bio-Budget) following the manufacturer’s instructions and a CFX96 Real-Time System machine (BioRad). See supplementary table 1 for primers.

### Electrophoretic light scattering measurements

To ensure homogeneous measurement conditions and best visibility of ζ-potential changes, vectors were submitted to buffer exchange prior to ELS measurement. Buffer exchange was performed using 5E10 VPs and PD-10 MiniTrap G-25 columns (GE Healthcare, Solingen, Germany) following the companies instructions with the “gravity” protocol. Briefly, the column was equilibrated using the measurement buffer (50 mM HEPES, pH 7.2). Afterwards, vectors were added to the column and subsequently, the vector volume was adjusted to 500 μL by adding measurement buffer to the column. The column was placed on an 1.5 mL reaction tube and the vector was eluted using 1 mL measurement buffer. Complete sample volume was used to measure “particle concentration” (Zetasizer Advance Serie – Ultra Red, Malvern Panalytical, Kassel, Germany) in a glass cuvette with square aperture (PCS1115). Thereby, size and concentration were determined by multiple angle dynamic light scattering (MADLS) with three measurement repeats (25°C, dispersant scattering mean count rate 179 kcps, dispersant values: R.I. 1.33; viscosity 0.8872 mPa s). For ζ-potential measurement, 700 μL of the suspension were transferred to a folded capillary cell (DTS1070) and measured as followed. To ensure sample integrity, three size measurements in backscatter mode where done before and after the ζ-potential measurement (25°C, dispersant values as given above). ζ-potential measurement was done in “general purpose” mode with a minimum number of runs of 10, three repeats and 60 seconds pause between each repeat (25°C, dispersant values as given above).

### Aggregate Pull-Down

1E7 VPs were incubated for 30 min. at 37°C in 30 μL of PBS + 1% BSA with or without 10 μg/mL of PF4. After centrifugation for 5 min. At 1000 g, the supernatant was transfered to a new tube containing 30 μL of 2x alkaline lysis buffer (50 mM NaOH + 4 mM EDTA), while 30 μL of 2x alkaline lysis buffer and 30 μL of PBS + 1% BSA were added to the pellet. Both treated supernatant and pellet were mixed thoroughly and heated at 95°C for 10 min. in order to release Ad genomes, then neutralized with 20 μL of cold neutralisation solution and titrated by qPCR (see supplementary table 1 for primers).

### Erythrocyte Pull-Down

This assay was inspired from (*18*).Venous blood was collected into EDTA tubes (Sarstedt, 02.1066.001) from the antecubital vein of a healthy volunteer who gave informed consent. The blood was swiveled at RT for 15 min. then centrifugated for 5 min. at 2000 g in order to isolate erythrocytes, which were then washed three times with PBS, resuspended in PBS + 1% BSA to a concentration of 5.5e9 cells / mL, and kept at 4°C for no more than 3 days. 4E7 VPs were incubated 30 min. at 37°C in 80μL of the erythrocyte suspension with or without 10 μg/mL of PF4. The suspension was centrifugated for 5 min. at 1000 g and the supernatant was transfered to a new tube. Both supernatant and pellet were treated with 2x alkaline lysis buffer, mixed thoroughly and heated at 95°C for 10 min. in order to release Ad genomes, then neutralized with neutralisation solution and titrated by qPCR (see supplementary table 1 for primers).

The study was approved by the ethics committee of the University Witten/Herdecke (approval number 216/2020, December 17^th^ 2020).

### Cell culture

The A549-ΔCAR and A549-ΔCD46 cell lines were constructed by deleting respectively the CAR and CD46 receptors using CRISPR-Cas9 (*19*). Cells were cultivated with Dulbecco’s Modified Eagle Medium (DMEM, Pan-Biotech; for 293, A549-WT, A549-ΔCAR and A549-ΔCD46 cells) or Roswell Park Memorial Institute 1640 Medium (RPMI, Pan-Biotech; for SKOV-3 cells), each supplemented with 10% Fetal Bovine Serum (FBS, Pan-Biotech) and Penicillin-Streptomycin (P/S, Pan-Biotech) at 37°C under an atmosphere with 5% CO_2_. Cells were tested for mycoplasma infection using the VenorGeM OneStep kit (Minerva Biolabs).

### Infectivity assays

To test the impact of PF4 on the infectivity of GLN library vectors in A549-derived cells, 2E7 VP/mL were incubated 10 min. at 37°C in OptiMEM (Gibco) with or without 10 μg/mL of PF4. At the end of incubation time, the culture medium of subconfluent cells was replaced by the virus suspension, counting 20 VP/cell (vpc). Part of the infected cells were kept to measure early Ad gene expression by luciferase assay or flow cytometry, while the others were washed three times and harvested 3 hours post infection (hpi) to titrate internalized Ad genomes.

To quantify early infection rates, cell DNA was extracted using the Monarch genomic DNA purification kit (NEB #T3010L) or the NucleoSpin Tissue kit (Macherey-Nagel #740952-250) following manufacturer’s instructions. Internalized Ad genomes and cell genomes and were titrated by qPCR and the number of Ad genomes per cell was used as an estimator of infectivity (see supplementary table 1 for primers).

Luciferase luminescence was measured 24 hpi using the Nano-Glo^®^ Luciferase Assay (Promega, Madison, USA) kit, a TECAN infinite f plex plate reader and black 96-well luciferase plates (Thermo Fisher Scientific Nunc A/S).

GFP fluorescence intensity was measured 24 hpi. Cells were harvested, washed twice in PBS, fixated 10 min. in 2% formaldehyde, washed twice in PBS then resuspended in PBS and analysed by flow cytometry (CytoFlex, Beckman Coulter, Munich, Germany) in FITC channel (585/42 nm), excited with a 488 nm laser.

The influence of PF4 on fX-mediated transduction efficiency of Ad5-ΔE1 into SKOV-3 cells was investigated by measurements of Ad5-ΔE1 genome uptake (qPCR) and eGFP expression (flow cytometry). SKOV-3 cells were seeded 24 h prior to transduction in 24-well-plates for qPCR (4E5 cells/well) or in 96-well-plates for flow cytometry (3E4 cells/well). For the following experiments, RPMI was supplemented with 400μM CaCl_2_. 1000 vpc were incubated in either 200 μL (qPCR) or 100 μL (flow cytometry) of RPMI, RPMI + 8 μg/mL fX, RPMI + 8 μg/mL PF4, or RPMI + 8 μg/mL PF4 and fX for 10 minutes at room temperature. To test whether sequential addition of the interaction partners might have an effect on fX-mediated Ad5-ΔE1 transduction, VPs were incubated in either 100 μL (qPCR) or 50 μL (flow cytometry) of RPMI + 16 μg/mL fX or 16 μg/mL PF4 for 10 minutes at room temperature. Subsequently, the same volume with the same concentration of the other interaction partner was added and SKOV-3 cell medium was replaced with the suspensions described above. Cells were incubated at 37°C and 5 % CO_2_.

For qPCR (24-well-plates), 3 h post transduction, cells were washed 5 times with warm PBS to remove extracellular VPs. Cells were detached with trypsin, centrifuged at 400 g for 6 minutes and resuspended in 200 μL PBS. Total genomic DNA was isolated as described in (*29*) using proteinase and RNAse treatment as well as phenol chloroform extraction. qPCR was performed with 2 μL of sample DNA (5 ng/μL). Ad5-ΔE1 genomes were titrated relative to the number of SKOV-3 genomes (see supplementary table 1 for primers). qPCR conditions were 2’ 50°C; 3’ 95°C; 41 cycles of 45’’ 95°C denaturation, 45’’ 60°C annealing and elongation. For flow cytometry (96-well-plates), 72 h post transduction, cells were washed three times with warm PBS to remove extracellular VPs. Cells were detached with trypsin, centrifuged at 400 g for 6 minutes and resuspended in 100 μL PBS. eGFP expression was analyzed via flow cytometry (CytoFlex, Beckman Coulter, Munich, Germany) in FITC channel (585/42 nm), excited with a 488 nm laser.

### Sequence and protein structure analyses

Alignments of hexon protein sequences were performed using the Mafft online tool with default parameters (*31*). Models of Ad14, Ad34 and Ad35 hexon monomer structures were constructed on SWISS-MODEL (*32*) with the default parameters, using RCSB Protein Data Bank’s 3tg7 structure of Ad5 hexon monomer as template (*33*). Graphic visualizations were performed with chimeraX (*34*).

### Statistical analyses

In ELISA-qPCR screens of Ad collections (figures 2a, 4b), usual statistical tests were irrelevant given that our goal was to identify vectors which do not bind to PF4, and not the opposite. Therefore, non-binding vectors were considered to be those for which the number of bound VPs in PF4-coated wells overlapped with that in non-coated wells in all experiment repeats (supplementary figure 2).

When N≤4 data points had been acquired, error bars indicate the minimum and maximum range.

When N≥5, error bars indicate standard deviation and pairwise comparisons were performed using Mann-Whitney U tests when applicable. Multiple comparisons were conducted with Welch’s ANOVA. In figures 4d and 4f, we tested whether PF4-associated fold increases were significantly different from 1 using one-sample Wilcoxon signed rank test. The significance threshold was set at p<0.05. Significance symbols: ns = non significant, * = p<0.05, ** = p<0.01, *** = p<0.001 Statistical analyses and visualizations were performed with the R software with the packages ggplot2 and dplyr (*35*, *36*).

