## supplementary figures and video for "Adenovirus type 34 and HVR1-deleted Adenovirus type 5 do not bind to PF4: clearing the path towards vectors without thrombosis risk"

### Slide 1
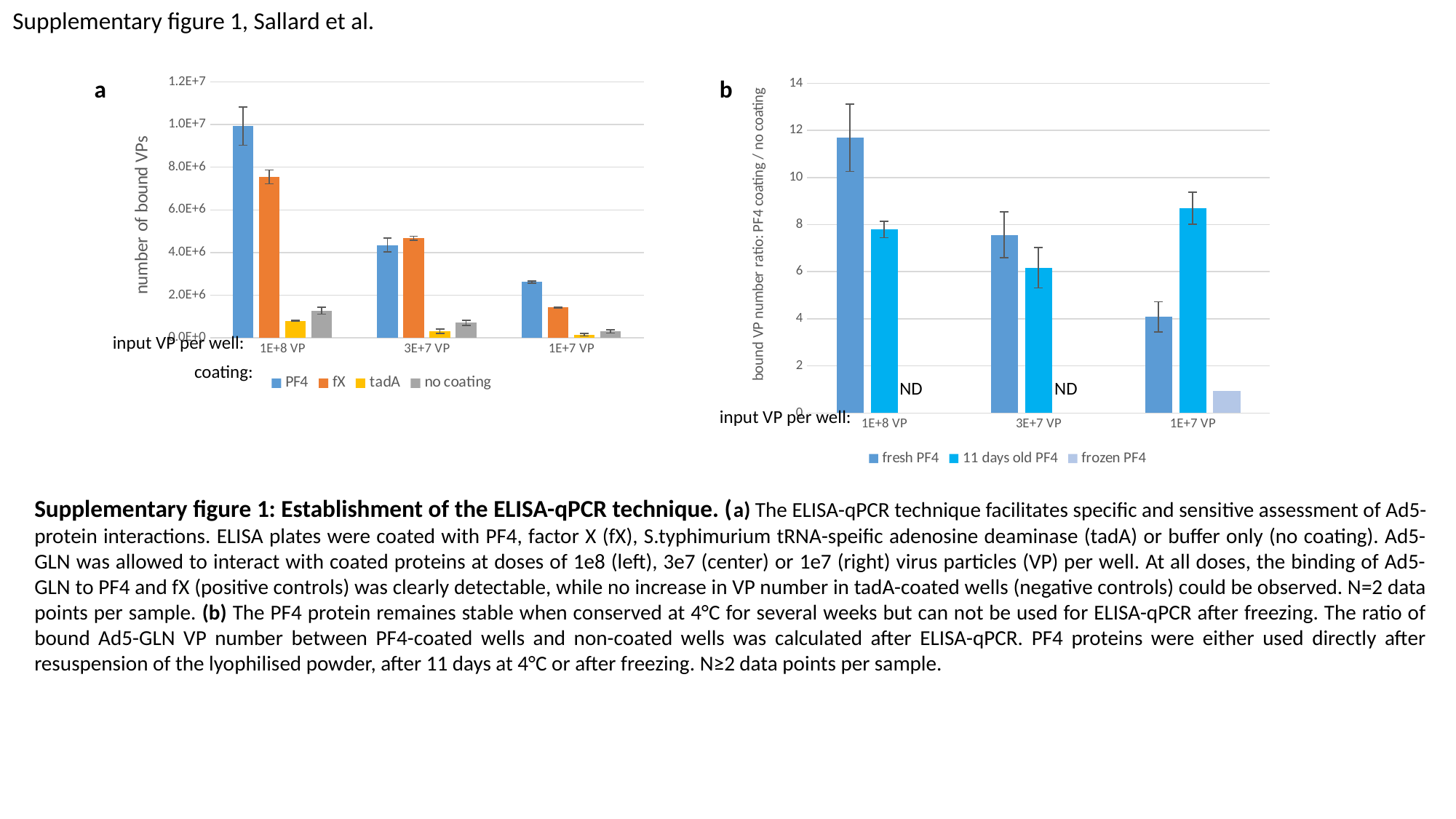

Supplementary figure 1, Sallard et al.
a
#### Chart
| Category | PF4 | fX | tadA | no coating |
|---|---|---|---|---|
| 1E+8 VP | 9927039.108151749 | 7544307.557110665 | 791157.846394926 | 1272888.6538057094 |
| 3E+7 VP | 4352576.884978228 | 4666009.000900851 | 306330.1319049131 | 705081.0380761642 |
| 1E+7 VP | 2618153.9663668997 | 1420193.8820868619 | 153779.4430717563 | 300999.6610282516 |input VP per well:
coating:
#### Chart
| Category | fresh PF4 | 11 days old PF4 | frozen PF4 |
|---|---|---|---|
| 1E+8 VP | 11.694642323909415 | 7.798827555318116 | None |
| 3E+7 VP | 7.570016812170659 | 6.173158332061193 | None |
| 1E+7 VP | 4.079985691817927 | 8.698195730264167 | 0.9382508583477875 |input VP per well:
ND	 ND
b
Supplementary figure 1: Establishment of the ELISA-qPCR technique. (a) The ELISA-qPCR technique facilitates specific and sensitive assessment of Ad5-protein interactions. ELISA plates were coated with PF4, factor X (fX), S.typhimurium tRNA-speific adenosine deaminase (tadA) or buffer only (no coating). Ad5-GLN was allowed to interact with coated proteins at doses of 1e8 (left), 3e7 (center) or 1e7 (right) virus particles (VP) per well. At all doses, the binding of Ad5-GLN to PF4 and fX (positive controls) was clearly detectable, while no increase in VP number in tadA-coated wells (negative controls) could be observed. N=2 data points per sample. (b) The PF4 protein remaines stable when conserved at 4°C for several weeks but can not be used for ELISA-qPCR after freezing. The ratio of bound Ad5-GLN VP number between PF4-coated wells and non-coated wells was calculated after ELISA-qPCR. PF4 proteins were either used directly after resuspension of the lyophilised powder, after 11 days at 4°C or after freezing. N≥2 data points per sample.

### Slide 2
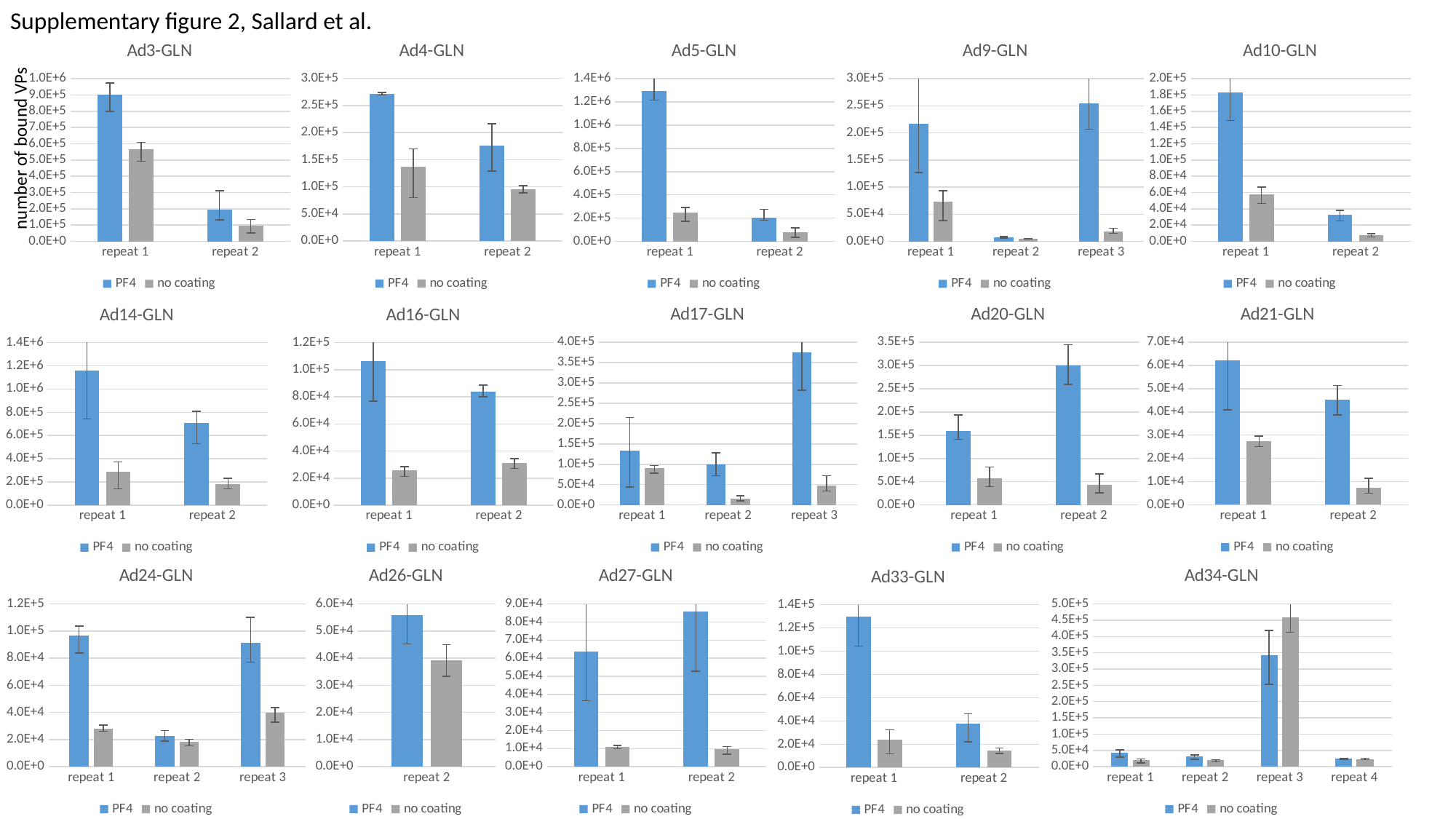

Supplementary figure 2, Sallard et al.
#### Chart: Ad4-GLN
| Category | PF4 | no coating |
|---|---|---|
| repeat 1 | 271827.04636852 | 137543.8266626262 |
| repeat 2 | 175763.5731971248 | 95191.54362631142 |
#### Chart: Ad3-GLN
| Category | PF4 | no coating |
|---|---|---|
| repeat 1 | 901449.0588860225 | 567442.9807324721 |
| repeat 2 | 196341.0633464315 | 94810.96188574347 |
#### Chart: Ad5-GLN
| Category | PF4 | no coating |
|---|---|---|
| repeat 1 | 1294030.6609899169 | 249715.6756789606 |
| repeat 2 | 202682.86831432252 | 77743.17063665169 |
#### Chart: Ad9-GLN
| Category | PF4 | no coating |
|---|---|---|
| repeat 1 | 216654.2462784504 | 73820.71776371663 |
| repeat 2 | 7721.8561289315985 | 4980.402066769913 |
| repeat 3 | 254073.80412414836 | 18728.252749476927 |
#### Chart: Ad10-GLN
| Category | PF4 | no coating |
|---|---|---|
| repeat 1 | 183047.41506719973 | 57967.5685583688 |
| repeat 2 | 33065.082876141365 | 7779.967484802968 |number of bound VPs
#### Chart: Ad21-GLN
| Category | PF4 | no coating |
|---|---|---|
| repeat 1 | 62238.518039323644 | 27294.086121869856 |
| repeat 2 | 45404.420485670584 | 7334.328366397982 |
#### Chart: Ad17-GLN
| Category | PF4 | no coating |
|---|---|---|
| repeat 1 | 133681.54112264203 | 90057.83014475174 |
| repeat 2 | 100253.44905216334 | 15867.896365004979 |
| repeat 3 | 374989.27502806945 | 47820.09693232124 |
#### Chart: Ad20-GLN
| Category | PF4 | no coating |
|---|---|---|
| repeat 1 | 158966.90983225763 | 56994.8790919251 |
| repeat 2 | 300455.91422380455 | 42616.973099337214 |
#### Chart: Ad16-GLN
| Category | PF4 | no coating |
|---|---|---|
| repeat 1 | 106527.53097325226 | 25619.323295001766 |
| repeat 2 | 83702.7031071722 | 31166.02779128622 |
#### Chart: Ad14-GLN
| Category | PF4 | no coating |
|---|---|---|
| repeat 1 | 1161630.8514969747 | 289890.4669991371 |
| repeat 2 | 705111.669610663 | 180062.1157840849 |
#### Chart: Ad34-GLN
| Category | PF4 | no coating |
|---|---|---|
| repeat 1 | 41951.17407632769 | 19845.328679202852 |
| repeat 2 | 31309.463299664523 | 19545.050463210417 |
| repeat 3 | 342943.9728711116 | 459807.0811569507 |
| repeat 4 | 24148.05069431715 | 23463.179315759662 |
#### Chart: Ad26-GLN
| Category | PF4 | no coating |
|---|---|---|
| repeat 2 | 55763.82109022635 | 39183.60652824 |
#### Chart: Ad27-GLN
| Category | PF4 | no coating |
|---|---|---|
| repeat 1 | 63805.53193032348 | 10765.653591622642 |
| repeat 2 | 85659.51264017704 | 9582.548994965655 |
#### Chart: Ad24-GLN
| Category | PF4 | no coating |
|---|---|---|
| repeat 1 | 96938.7065627284 | 27832.42643685302 |
| repeat 2 | 22676.58617761156 | 18436.632858910027 |
| repeat 3 | 91406.43959909033 | 39691.6004511885 |
#### Chart: Ad33-GLN
| Category | PF4 | no coating |
|---|---|---|
| repeat 1 | 129889.26204029676 | 23754.107061611427 |
| repeat 2 | 37515.81092861296 | 14311.410091616766 |

### Slide 3
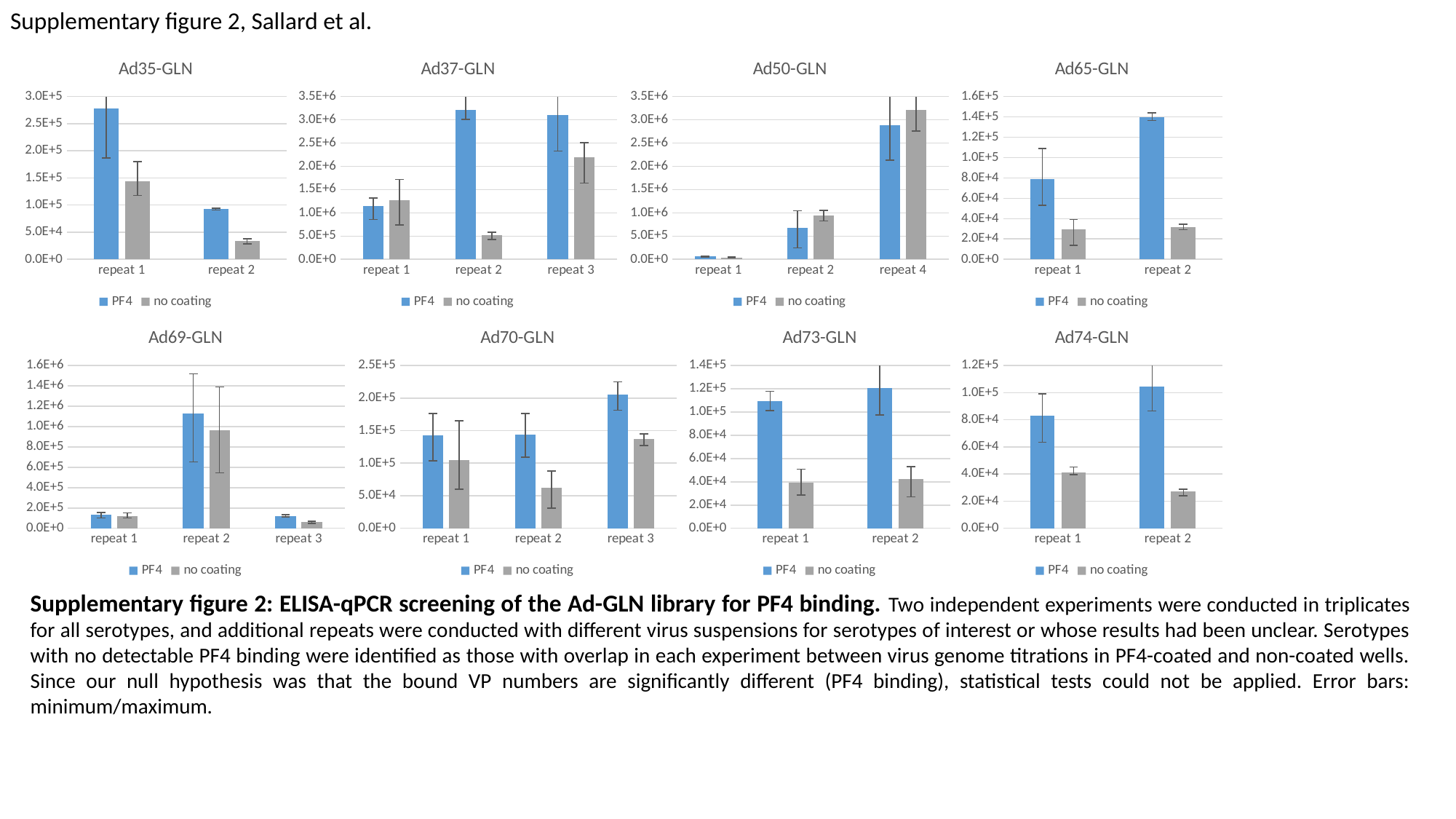

Supplementary figure 2, Sallard et al.
#### Chart: Ad50-GLN
| Category | PF4 | no coating |
|---|---|---|
| repeat 1 | 60311.1092372229 | 40824.13589470409 |
| repeat 2 | 682775.2529994077 | 939958.2753274683 |
| repeat 4 | 2876958.3585958555 | 3215675.842668672 |
#### Chart: Ad37-GLN
| Category | PF4 | no coating |
|---|---|---|
| repeat 1 | 1148546.7619702125 | 1264851.7020648245 |
| repeat 2 | 3218777.4580089487 | 512768.92660171614 |
| repeat 3 | 3105983.2660111827 | 2191774.325047259 |
#### Chart: Ad35-GLN
| Category | PF4 | no coating |
|---|---|---|
| repeat 1 | 277892.25554577634 | 144234.73852224645 |
| repeat 2 | 92826.51255646604 | 34210.90721594056 |
#### Chart: Ad65-GLN
| Category | PF4 | no coating |
|---|---|---|
| repeat 1 | 79011.68185342554 | 29584.561238852963 |
| repeat 2 | 139424.93427836607 | 31934.643786158365 |
#### Chart: Ad74-GLN
| Category | PF4 | no coating |
|---|---|---|
| repeat 1 | 82881.383044747 | 41415.45393678273 |
| repeat 2 | 104448.16323087893 | 27046.161276185492 |
#### Chart: Ad73-GLN
| Category | PF4 | no coating |
|---|---|---|
| repeat 1 | 109310.8234784154 | 39437.84203597935 |
| repeat 2 | 120489.75543780094 | 42137.88156458884 |
#### Chart: Ad70-GLN
| Category | PF4 | no coating |
|---|---|---|
| repeat 1 | 143020.85392339397 | 104659.64074598088 |
| repeat 2 | 143695.71993907972 | 62208.156246886734 |
| repeat 3 | 205417.93885882356 | 136727.62202981635 |
#### Chart: Ad69-GLN
| Category | PF4 | no coating |
|---|---|---|
| repeat 1 | 135545.11956317964 | 122840.99659494233 |
| repeat 2 | 1128183.0461662125 | 963460.9102138799 |
| repeat 3 | 119604.90525209701 | 59276.2546515116 |Supplementary figure 2: ELISA-qPCR screening of the Ad-GLN library for PF4 binding. Two independent experiments were conducted in triplicates for all serotypes, and additional repeats were conducted with different virus suspensions for serotypes of interest or whose results had been unclear. Serotypes with no detectable PF4 binding were identified as those with overlap in each experiment between virus genome titrations in PF4-coated and non-coated wells. Since our null hypothesis was that the bound VP numbers are significantly different (PF4 binding), statistical tests could not be applied. Error bars: minimum/maximum.

### Slide 4
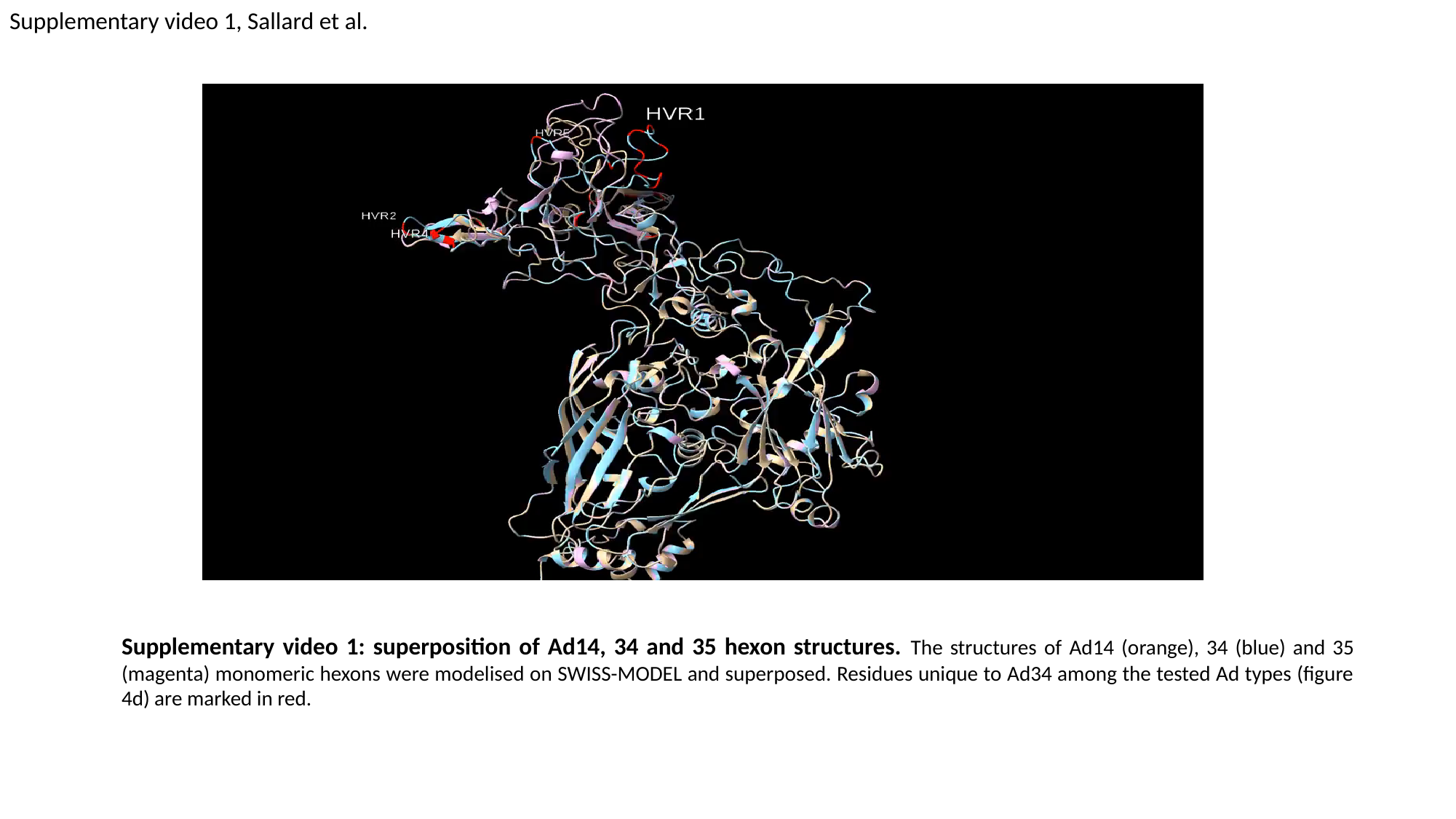

Supplementary video 1, Sallard et al.
Supplementary video 1: superposition of Ad14, 34 and 35 hexon structures. The structures of Ad14 (orange), 34 (blue) and 35 (magenta) monomeric hexons were modelised on SWISS-MODEL and superposed. Residues unique to Ad34 among the tested Ad types (figure 4d) are marked in red.
